# Functional MRI of native and non-native speech production in sequential German-English bilinguals

**DOI:** 10.1101/2021.02.18.431784

**Authors:** Miriam Treutler, Peter Sörös

## Abstract

Bilingualism and multilingualism are highly prevalent. Non-invasive brain imaging has been used to study the neural correlates of native (L1) and non-native (L2) speech and language production, mainly on the lexical and syntactic level. Here, we acquired continuous fast event-related FMRI during visually cued overt production of exclusively German and English vowels and syllables. We analyzed data from 13 university students, native speakers of German and sequential English bilinguals. The production of non-native English sounds was associated with increased activity of the left primary sensori-motor cortex, bilateral cerebellar hemispheres (lobule VI), left inferior frontal gyrus, and left anterior insula compared to native German sounds. The contrast German > English sounds was not statistically significant. Our results emphasize that the production of non-native speech requires additional neural resources already on a basic phonological level in sequential bilinguals.

## Introduction

Bilingualism and multilingualism, the ability to communicate in two or more languages, are highly prevalent. Although an exact definition of bilingualism and precise statistics are missing, it is estimated that more than 50% of the global population actively use more than one language (1). At least 55 countries have two or more official languages.^1^ Many individuals are exposed to and use two languages on a daily basis from birth or starting in their first years of life (simultaneous or early bilinguals). Many others learn at least one foreign language (L2) at school or later in life (sequential or late bi- or multilinguals). In the European Union, 95% of all students in upper secondary education learn English as a foreign language, 22% Spanish, 18% French, and 17% German.^2^

With the advent of non-invasive methods of brain research, such as event-related potentials (ERPs), positron emission tomography (PET), and functional MRI (FMRI), the neural correlates of bilingual speech and language production became readily accessible to scientific research, contributing to the extensive increase of published studies on bilingualism in the past two decades (2). Numerous studies demonstrated that L2 production relies on neural systems that are also used in monolinguals, with often increased brain activity for L2 production due to cross-linguistic interference during lexical retrieval, articulatory planning, articulation, and auditory and sensory feedback (3). Most work on the organization of the bilingual brain has been performed on the lexical and syntactic level (4, 5). Considerably less research has been done on bilingual speech motor control (6).

We have previously investigated the production of speech sounds of different complexity frequently used in the participants’ native language with clustered FMRI acquisition (or sparse sampling) (7, 8). We found that the production of an isolated vowel (“a”), a consonant-vowel syllable (“pa”, “ta”, or “ka”), and a trisyllabic utterance (“pataka”) was associated with the activation of a distributed neural network of cortical and subcortical brain regions, including the primary sensorimotor cortex, the supplementary motor area, the cerebellum, and the superior temporal gyrus. The production of the more complex “pataka”, as compared to “a”, resulted in increased activity in the left inferior frontal gyrus, the left cerebellar hemisphere, and the bilateral temporal cortex (7).

In the present study, we investigated the production of speech sounds commonly used in German (but unknown in English) and speech sounds commonly used in English (but unknown in German) in late bilingual German university students who grew up in a monolingual German-speaking family and started to learn English at school. We used, in contrast to our earlier studies (7, 8), continuous fast event-related FMRI, after pilot measurements demonstrated moderate head motion during overt speech production, corroborating the results of a recent FMRI study on overt sentence production (9). We hypothesized that production of non-native speech sounds should resemble the production of native, more complex sounds, i.e. should be associated with increased activity in key areas of speech motor control (such as the left inferior frontal gyrus and the cerebellar hemispheres).

## Methods

### Participants

For the present study, 15 healthy young adults were investigated. As two participants had to be excluded because of incorrect task performance (see section *Behavioral data analysis*), the following data analyses are based on 13 participants (7 women, 6 men) with a mean age ± standard deviation of 25.5 ± 3.0 years (minimum: 20 years, maximum: 32 years). All participants were native speakers of German (native language, L1) and started to learn English at school after the age of 6 years (first foreign language, L2). Participants self-rated their English proficiency between the levels B1/B2 and C1, according to the Common European Framework of Reference for Languages.^3^ B1 is considered intermediate, B2 upper intermediate, and C1 advanced proficiency of a foreign language. According to the Edinburgh Handedness Inventory–Short Form (10), nine participants were right-handed (handedness scores: 62.5 - 100) and 4 participants were bimanual (handedness score: 50).

This experiment was part of a larger project on oral and speech language functions. Detailed inclusion and exclusion criteria for the participants were published in a paper on the neural correlates of tongue movements (11). In brief, all participants were part of a convenience sample of students of the University of Oldenburg, Germany, without a history of neurological or psychiatric disorders or substance abuse. All participants gave written informed consent for participation in the study. A compensation of 10 C per hour was provided. The study was approved by the Medical Research Ethics Board, University of Oldenburg, Germany (2017-072).

### Paradigm

During the experiment, participants were visually cued to articulate one of the following four vowels or syllables (the symbols of the International Phonetic Alphabet^4^ are given in brackets): “ö” (ø:), “aw” (ɔ:), “che” (çə), and “the” (ðə). The vowel “ö” and the syllable “che” are common in German, but do not exist in standard English. By contrast, the vowel “aw” and the syllable “the” are common in English, but do not exist in German. Especially the English “th” (ð) is notoriously difficult to pronounce for Germans and is usually spoken with a characteristic German accent.

The corresponding letters were projected onto a screen with an LCD projector and presented to the participants in the scanner through a mirror on the head coil using Cogent 2000 v125^5^ run in MATLAB R2015b.

Using a fast event-related design, 120 visual stimuli (30 per condition) were shown in a pseudorandomized order for 1000 ms. During the interstimululs interval, a fixation cross was presented for a variable length of time, between 2000 and 8000 ms. The measurement started with the presentation of a fixation cross for 5000 ms and ended with the presentation of a fixation cross for another 15000 ms.

Before the FMRI experiment, a short training run was presented on the stimulation PC outside the scanner to give all participants the opportunity to familiarize themselves with the paradigm. All participants were instructed to articulate the corresponding sounds as soon as the letters appeared on the screen in the loudness of a regular conversation and to keep their head as still as possible.

After the experiment described here, three additional experiments were performed during the same imaging session, including the overt production of tongue twisters, movements of the tongue (11), and overt production of sentences^6^. The duration of the entire scanning session was approximately 45 min.

### MR data acquisition

Structural and functional MR images of the entire brain were acquired on a research-only Siemens MAGNETOM Prisma whole-body scanner at 3 Tesla (Siemens, Erlangen, Germany) and a 64-channel head/neck receive-array coil located at the Neuroimaging Unit, School of Medicine and Health Sciences, University of Oldenburg, Germany.^7^ We used a T1-weighted MPRAGE sequence to acquire structural data and a T2*-weighted BOLD sequence (305 volumes, time of acquisition: 9:16 min) to acquire functional data (for details, see (11)).

### Audio recording

During the FMRI experiment, all utterances were recorded on a PC through an FMRI-compatible microphone attached at the head coil (FOM1-MR, Micro Optics Technologies, Middleton, WI, USA).

### Behavioral data analysis

Relative (volume-to-volume) and absolute (relative to the middle volume) head motion were determined by volume-realignment using MCFLIRT (FSL version 6.00) (12).

All audio recordings were checked for correct task performance. One participant misunderstood the task and spoke all sounds twice, another participant pronounced all sounds considerably longer than demonstrated during the pre-scan training. Both participants were excluded from the further data analysis. The remaining 13 participants produced four wrong or unintelligible sounds (coughing); these individual sounds were also excluded from the FMRI data analysis. Because of the continuous gradient noise during scanning, further analyses (e.g. speech latencies or phonetic analyses) were not possible.

### Preprocessing of functional images

MRI analyses were done on University of Oldenburg’s high-performance computer cluster CARL. Preprocessing was performed using fM-RIPrep version 20.0.5 (RRID:SCR_016216) (13). For a detailed description see^8^. Functional data were motion corrected by volume-realignment using MCFLIRT and registered to the MNI152NLin6Asym standard space template. ICA-based Automatic Removal Of Motion Artifacts (AROMA)^9^ was used to denoise the functional images, using the non-aggressive option (14). Slicetime correction was not performed. Preprocessing reports for all participants are available at the Open Science Framework (OSF).^10^

### First-level FMRI analysis

Preprocessed functional data sets were analyzed with FEAT (FSL version 6.00), performing a general linear model-based time-series analysis using voxel-wise multiple linear regressions (15).

The time courses of the two German sounds and the two English sounds were convolved with a gamma hemodynamic response function (phase: 0 s, standard deviation: 3 s, mean lag: 6 s) and served as regressors of interest. The temporal derivative of each primary regressor was included as a regressor of no interest to improve the model fit to account for differences in response latency. Regressors of interest (experimental conditions) and regressors of no interest (temporal derivatives) formed the design matrix used for voxel-wise multiple linear regressions. Motion parameters and physiological noise regressors were not included in the design matrix because ICA-AROMA was used for denoising. To remove temporal autocorrelations, time-series pre-whitening was used (16). After generating parameter estimates (PEs) for every primary regressor and every participant, the following contrasts of parameter estimates (COPEs) were calculated: 1) German > rest, 2) English > rest, 3) German > English, and 4) English > German. Z statistic images were thresholded non-parametrically using a cluster-forming threshold of Z > 2.3 and a (corrected) cluster significance threshold of p < 0.05. Brain activity maps of all 13 participants are available at OSF.

### Second-level FMRI analysis

Mixed-effects group analysis maps were generated by FLAME (stages 1 and 2) for all contrasts. Again, Z statistic images were thresholded at Z > 2.3 (p < 0.05). Brain activity maps for all contrasts are available at OSF.

Local maxima (peaks of brain activity) were identified within the Z statistic images using FSL’s cluster command (minimum distance between local maxima: 10 mm; 62 local maxima were found). The anatomical location of each local maximum was determined with FSL’s atlasquery command and the following probabilistic atlases (for details see^11^): 1) Harvard-Oxford cortical structural atlas (48 cortical areas), n2) Harvard-Oxford subcortical structural atlas (21 subcortical areas), and 3) Probabilistic cerebellar atlas (28 regions) (17). The local maximum with the highest Z value in a given anatomical area is reported in **Table 1**. Because we report local maxima of brain activity, the extent of activity (cluster size) cannot be determined.

**Table 1.** Local maxima of brain activity: stereotaxic coordinates in MNI space, Z values, and corresponding brain regions for the contrast frontal English (L2) > German (L1).

| Region | x (mm) | y (mm) | z (mm) | Z value |
| --- | --- | --- | --- | --- |
| L precentral gyrus | -46 | -12 | 32 | 3.24 |
| L postcentral gyrus | -60 | -6 | 28 | 4.37 |
| L inferior frontal gyrus (triangular) | -48 | 26 | 4 | 3.70 |
| L inferior frontal gyrus (opercular) | -54 | 14 | 26 | 3.53 |
| L insula | -28 | 22 | 0 | 3.50 |
| L cerebellum Lobule VI | -16 | -58 | -24 | 3.12 |
| R cerebellum Lobule VI | 14 | -68 | -28 | 3.59 |
| L cuneus | -4 | -86 | 22 | 4.75 |
| R cuneus | 6 | -76 | 28 | 4.29 |
| L intracalcarine cortex | -6 | -80 | 6 | 3.98 |
| R intracalcarine cortex | 10 | -76 | 16 | 4.65 |
| L occipital pole | -8 | -98 | -2 | 4.61 |
| R occipital pole | 18 | -88 | -2 | 4.70 |
| L lingual gyrus | -10 | -88 | -10 | 4.17 |
| R lingual gyrus | 4 | -74 | 0 | 3.87 |
| L occipital fusiform gyrus | -14 | -82 | -20 | 3.69 |
| R occipital fusiform gyrus | 24 | -72 | -6 | 3.76 |

## Results

### Head motion

**Figure 1** displays the relative displacement between two adjacent MRI volumes for all participants. In one participant, three values > 0.5 mm were found (1.38, 0.95, and 0.51 mm). In another participant, one value was 0.65 mm. All other values were less than 0.5 mm. The median relative displacement for all participants and all time-points was 0.07 mm. The maximum absolute displacement between the middle volume as a reference and all other volumes was less than 3 mm in all participants (minimum: 0.26 mm, maximum: 2.89 mm).

**Fig 1.**
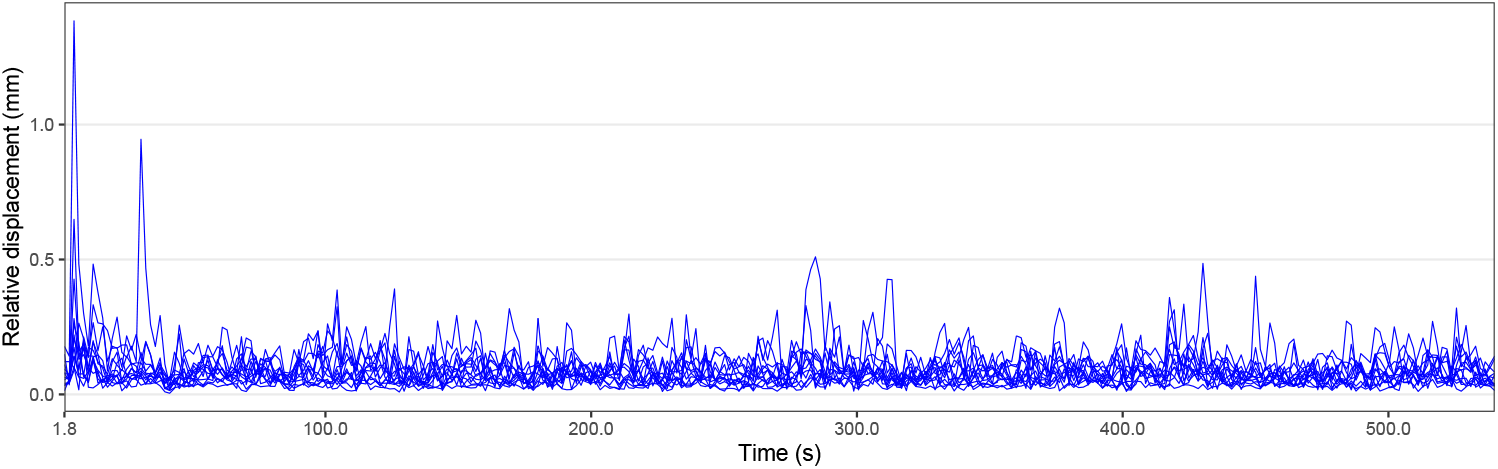
Individual head motion over time. Relative displacement, the distance between one volume and the following volume, during all FMRI measurements is displayed in blue.

### Brain activity

Production of German (L1) and English (L2) speech sounds, compared to baseline, was associated with similar and widespread activation of cortical and subcortical areas, primarily related to speech motor control, phonological processing, and visual processing (data shown at OSF). **Figure 2** illustrates brain areas significantly more active during production of English compared to German sounds. These areas include key regions of speech motor control (left lateral sensorimotor cortex, left inferior frontal gyrus, left anterior insula, and bilateral cerebellar hemispheres). In addition, Figure 2 displays brain areas more active during visual processing of English compared to German cues, including the bilateral lingual gyrus and the bilateral occipital fusiform gyrus. **Table 1** summarizes the coordinates of local maxima in MNI space and the respective Z value. The reverse contrast, German > English, did not result in significant differences.

**Fig 2.**
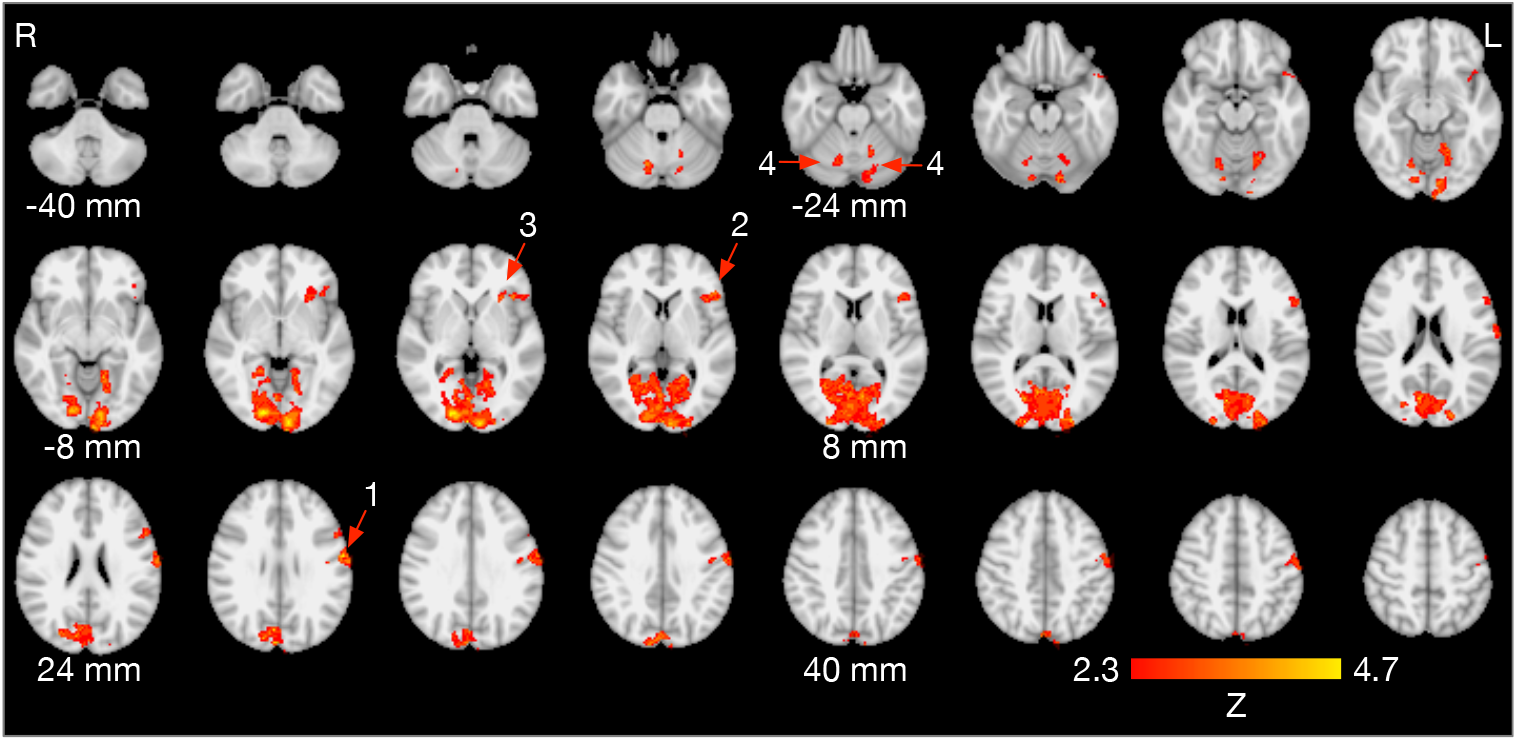
Brain activity for the contrast English > German speech production. The arrows point at the left lateral sensori-motor cortex (1), left inferior frontal gyrus (2), left anterior insula (3), and bilateral cerebellar lobule VI (4). Images are in radiological convention (the left hemisphere is seen on the right).

## Discussion

The present fast event-related FMRI study on the overt production of German and English speech sounds in native speakers of German, also proficient in English, demonstrated increased activity during non-native speech in critical areas of speech motor control. As speech production was cued by written letters, increased activity was also found in several occipital areas of the visual system (**Table 1, Figure 2**). Speech production is a highly complex task, depending on several integrated processing stages. During speech production, the brain rapidly retrieves phonological information, executes speech motor programs, encoding movement trajectories of the articulators, and monitors continuously auditory and somatosensory feedback. These processing stages are materialized in a widespread articulatory brain network, including key areas of the pyramidal and non-pyramidal motor system, and in a phonological network, primarily located in the temporal lobes (18, 19).

### Areas of increased brain activity

In the present study, production of non-native English sounds was associated with increased activity of the left primary sensori-motor cortex, bilateral cerebellar hemispheres (lobule VI), left inferior frontal gyrus, and left anterior insula.

The lateral primary sensori-motor cortex directly controls the muscles of the larynx (20, 21) and the articulators, including the tongue (11), and processes somatosensory information of the oral cavity (22). Although the laryngeal and oro-facial midline muscles are innervated by both hemispheres, specific speech motor plans, or articulatory gestures, are primarily represented in the left primary motor cortex (23). The primary motor cortex is not only involved in speech production, but also in speech perception, presumably encoding distinctive phonetic features of individual speech sounds (24). The cerebellar hemispheres receive afferents from the primary motor cortex via the cortico-ponto-cerebellar tracts and support sensorimotor control and coordination of laryngeal, orofacial, and respiratory movements (25). Generally considered to be heavily engaged in the rapid sequencing of speech sounds, forming syllables and words as well as producing the rhythm and intonation of continuous speech (i.e., prosody) (25), the bilateral cerebellar hemispheres are also involved in the production of single vowels (7). Multiple lines of evidence suggest that the cerebellar hemispheres are organized in a homuncular topology. Electric stimulation during neurosurgery demonstrated that the movements of the face and mouth are primarily represented in the hemispheric lobule VI (26). A graph theoretical analysis further elucidated the critical role of hemispheric lobule VI in the speech production network (21).

The integrity of the left inferior frontal gyrus, although not part of the core motor system, has been linked to speech production since Broca’s seminal observations (27, 28). The triangular and the opercular part of the left inferior frontal gyrus and, based on recent cytoarchitectonic and receptorar-chitectonic analyses, adjacent frontal regions are the structural correlates of Broca’s area (29). In addition to its critical role in the left-hemispheric language network, Broca’s area is also believed to be part of the articulatory network (30). Direct cortical surface recordings in neurosurgical patients suggested that Broca’s area mediates the information flow between the temporal cortex, the likely anatomical substrate of phonological planning, and the primary motor cortex, thus preparing an appropriate articulatory code to be executed by the motor cortex (31). Deactivation of Broca’s area is associated with slowing of speech production (32, 33).

The insulae are areas of sensory, motor, cognitive, and affective integration, e.g. processing somatosensory (22, 34) and nociceptive information (35). The insulae are also involved in movements such as breathing (36), swallowing (37), and speech production (38, 39). The exact functions of the insulae in the articulatory network are under debate and still not entirely clear (40).

This study was designed to investigate the articulatory and phonological networks underlying L2 production. As we presented letters to cue verbal responses, we also found activity in parts of the visual system. We saw increased activity for the letter strings “the” and “aw” in the fusiform and lingual gyri, areas involved in letter recognition and orthographic to phonological mapping (41). Similarly, sentence reading in sequential bilinguals was associated with increased activity in the left fusiform gyrus when reading L2 compared to L1 (42).

### Mechanisms of increased brain activity

Several studies compared speech motor control during the production of L1 utterances of different complexities (7, 43–45) and found increased activity of the areas discussed above. In the present study, however, the formal complexity of the produced speech sounds was identical in German and English (a single vowel and a consonant-vowel syllable each). The interpretation of our results is not straightforward because our participants were late multilinguals, less proficient in English, and used English considerably less than German.

Focusing on proficiency and use, we may argue that the production of L2 speech sounds requires more resources on different levels of the articulatory network, because L2 production is not as over-learned as L1 and performed in a less automatic fashion. This explanation would lead us to predict that intense training of the required sounds would result in decreased activity in the articulatory network. This interpretation appears to be supported by Ghazi-Saidi et al., who trained native speakers of Spanish to pronounce French cognates (phonologically and semantically similar words across languages) in a native accent for 4 weeks (46). In a picture naming paradigm, the authors found increased activity for the contrast L2 > L1 only in a small area of the left insula, but not in other areas of the articulatory network.

Focusing on age of acquisition, by contrast, we may argue that our participants started to learn English when German speech production was already consolidated and deeply encoded in the articulatory network, resulting in less efficient articulatory gestures for English speech production after the maturation of the articulatory network. This notion would lead us to predict that simultaneous bilinguals should not differ in brain activity when producing speech sounds in one of their languages. The notion of a sensitive period for speech motor control is corroborated by a study on sentence reading in simultaneous and sequential bilinguals, all using both languages on a daily basis (42). While brain activity was similar for simultaneous bilinguals, sequential bilinguals demonstrated increased activity in the left inferior frontal gyrus and left premotor cortex when reading aloud in L2 compared to L1. Importantly, activity in these areas showed a significant positive correlation with age of acquisition.

### Foreign accent

The results of the present study may help to better understand the neural correlates of foreign accent. While simultaneous bilinguals usually speak in a native or native-like accent in their languages, most sequential bilinguals speak L2 with a foreign accent, even if they perform similar to natives on the lexical and grammatical level (47, p. 2). A foreign accent is characterized by deviations in pronunciation compared to the norms of native speech (48, p. 253), mostly due to phonetic and phonological mismatches between the native language (L1) and L2 and caused by interference or transfer of pronunciation rules (49, p. 177). Our results imply that, for sequential bilinguals, the neural correlates of L2 production differ from L1 already at the fundamental level of vowel and syllable production and emphasize why it is so difficult, and often impossible, to loose a foreign accent.

## ACKNOWLEDGEMENTS

The authors wish to thank Katharina Grote and Gülsen Yanç for assisting with MRI data acquisition as well as Stefan Harfst and Fynn Schwietzer, Scientific Computing, University of Oldenburg, Germany, for continuous support with the HPC cluster. The authors also appreciate the work of Sarah Schäfer, Sarah Schumacher, and Zoe Tromberend, who performed the other experiments for this project.

This study was supported by the Neuroimaging Unit, University of Oldenburg, funded by grants from the German Research Foundation (DFG; 3T MRI INST 184/152-1 FUGG and MEG INST 184/148-1 FUGG). The high-performance computer cluster CARL, University of Oldenburg, is funded by a grant from the German Research Foundation (DFG; INST 184/157-1 FUGG).

## Footnotes

1 https://www.uottawa.ca/clmc/55-bilingual-countries-world

2 https://ec.europa.eu/eurostat

3 https://www.efset.or

4 https://www.internationalphoneticalphabet.org/ipa-sounds/ipa-chart-with-sounds/

5 http://www.vislab.ucl.ac.uk/cogent.php

6 https://f1000research.com/posters/8-687

7 https://uol.de/en/medicine/biomedicum/neuroimaging-unit

8 https://fmriprep.org

9 https://github.com/maartenmennes/ICA-AROMA

10 https://osf.io/t9qcw/

11 https://fsl.fmrib.ox.ac.uk/fsl/fslwiki/Atlases

## References

1. E Bialystok, FI Craik, and G Luk. Bilingualism: consequences for mind and brain. Trends Cogn Sci, 16:240–250, 2012.

2. JF Kroll and CA Navarro-Torres. Bilingualism. In JT Wixted, editor, Stevens’ Handbook of Experimental Psychology and Cognitive Neuroscience. Wiley, Hoboken, NJ, USA, 4th edition, 2018.

3. N Del Maschio and J Abutalebi. Language organization in the bilingual and multilingual brain. In JW Schwieter, editor, The Handbook of the Neuroscience of Multilingualism, pages 199–213. Wiley, Hoboken, NJ, USA, 2019.

4. L Sabourin. fMRI research on the bilingual brain. Annu Rev Appl Linguist, 34:1–14, 2014.

5. JF Kroll, PE Dussias, K Bice, and L Perrotti. Bilingualism, mind, and brain. Annu Rev Linguist, 1:377–394, 2015.

6. AJ Simmonds, RJ Wise, and R Leech. Two tongues, one brain: imaging bilingual speech production. Front Psychol, 2:166, 2011.

7. P Sörös, LG Sokoloff, A Bose, AR McIntosh, SJ Graham, and DT Stuss. Clustered functional MRI of overt speech production. Neuroimage, 32:376–387, 2006.

8. P Sörös, A Bose, LG Sokoloff, SJ Graham, and DT Stuss. Age-related changes in the functional neuroanatomy of overt speech production. Neurobiol Aging, 32:1505–1513, 2011.

9. DH Berro, J. Lemée, LM Leiber, E Emery, P Menei, and A Ter Minassian. Overt speech feasibility using continuous functional magnetic resonance imaging: Isolation of areas involved in phonology and prosody. J Neurosci Res, 98:2554–2565, 2020.

10. JF Veale. Edinburgh Handedness Inventory - Short Form: a revised version based on confirmatory factor analysis. Laterality, 19:164–177, 2014.

11. P Sörös, S Schäfer, and K Witt. Model-based and model-free analyses of the neural correlates of tongue movements. Front Neurosci, 14:226, 2020.

12. M Jenkinson, P Bannister, M Brady, and S Smith. Improved optimization for the robust and accurate linear registration and motion correction of brain images. Neuroimage, 17: 825–841, 2002.

13. O Esteban, CJ Markiewicz, RW Blair, CA Moodie, AI Isik, A Erramuzpe, JD Kent, M Goncalves, E DuPre, M Snyder, H Oya, SS Ghosh, J Wright, J Durnez, RA Poldrack, and KJ Gorgolewski. fMRIPrep: a robust preprocessing pipeline for functional MRI. Nat Methods, 16:111–116, 2019.

14. RHR Pruim, M Mennes, D van Rooij, A Llera, JK Buitelaar, and CF Beckmann. ICA-AROMA: A robust ICA-based strategy for removing motion artifacts from fMRI data. Neuroimage, 112: 267–277, 2015.

15. KJ Friston, AP Holmes, KJ Worsley, J-P Poline, CD Frith, and RSJ Frackowiak. Statistical parametric maps in functional imaging: a general linear approach. Hum Brain Mapp, 2: 189–210, 1995.

16. MW Woolrich, BD Ripley, M Brady, and SM Smith. Temporal autocorrelation in univariate linear modeling of FMRI data. Neuroimage, 14:1370–1386, 2001.

17. J Diedrichsen, JH Balsters, J Flavell, E Cussans, and N Ramnani. A probabilistic MR atlas of the human cerebellum. Neuroimage, 46:39–46, 2009.

18. E Golfinopoulos, JA Tourville, and FH Guenther. The integration of large-scale neural network modeling and functional brain imaging in speech motor control. Neuroimage, 52: 862–874, 2010.

19. G Hickok. The architecture of speech production and the role of the phoneme in speech processing. Lang Cogn Process, 29:2–20, 2014.

20. S Brown, AR Laird, PQ Pfordresher, SM Thelen, P Turkeltaub, and M Liotti. The somatotopy of speech: phonation and articulation in the human motor cortex. Brain Cogn, 70:31–41, 2009.

21. K Simonyan and S Fuertinger. Speech networks at rest and in action: interactions between functional brain networks controlling speech production. J Neurophysiol, 113:2967–2978, 2015.

22. P Sörös, E Lalone, R Smith, T Stevens, J Theurer, RS Menon, and RE Martin. Functional mri of oropharyngeal air-pulse stimulation. Neuroscience, 153:1300–1308, 2008.

23. NE Neef, TN Hoang, A Neef, W Paulus, and M Sommer. Speech dynamics are coded in the left motor cortex in fluent speakers but not in adults who stutter. Brain, 138:712–725, 2015.

24. F Pulvermüller, M Huss, F Kherif, F Moscoso del Prado Martin, O Hauk, and Y Shtyrov. Motor cortex maps articulatory features of speech sounds. Proc Natl Acad Sci U S A, 103: 7865–7870, 2006.

25. H Ackermann. Cerebellar contributions to speech production and speech perception: psycholinguistic and neurobiological perspectives. Trends Neurosci, 31:265–272, 2008.

26. C Mottolese, N Richard, S Harquel, A Szathmari, A Sirigu, and M Desmurget. Mapping motor representations in the human cerebellum. Brain, 136:330–342, 2013.

27. PP Broca. Perte de la parole, ramollissement chronique et destruction partielle du lobe antérieur gauche. Bulletin de la Société d’Anthropologie, 2:235–238, 1861.

28. NF Dronkers, O Plaisant, MT Iba-Zizen, and EA Cabanis. Paul Broca’s historic cases: high resolution MR imaging of the brains of Leborgne and Lelong. Brain, 130:1432–1441, 2007.

29. K Zilles and K Amunts. Cytoarchitectonic and receptorarchitectonic organization in Broca’s region and surrounding cortex. Curr Opin Behav Sci, 21:93–105, 2018.

30. E Fedorenko and IA Blank. Broca’s area is not a natural kind. Trends Cogn Sci, 24(4): 270–284, 2020.

31. A Flinker, A Korzeniewska, AY Shestyuk, PJ Franaszczuk, NF Dronkers, RT Knight, and NE Crone. Redefining the role of Broca’s area in speech. Proc Natl Acad Sci U S A, 112: 2871–2875, 2015.

32. MA Long, KA Katlowitz, MA Svirsky, RC Clary, TM Byun, N Majaj, H Oya, MA Howard, and JDW Greenlee. Functional segregation of cortical regions underlying speech timing and articulation. Neuron, 89:1187–1193, 2016.

33. MK Leonard, M Desai, D Hungate, R Cai, NS Singhal, RC Knowlton, and EF Chang. Direct cortical stimulation of inferior frontal cortex disrupts both speech and music production in highly trained musicians. Cogn Neuropsychol, 36:158–166, 2019.

34. M Pugnaghi, S Meletti, L Castana, S Francione, L Nobili, R Mai, and L Tassi. Features of somatosensory manifestations induced by intracranial electrical stimulations of the human insula. Clin Neurophysiol, 122:2049–2058, 2011.

35. S Fazeli and C Büchel. Pain-related expectation and prediction error signals in the anterior insula are not related to aversiveness. J Neurosci, 38(29):6461–6474, 2018.

36. JL Herrero, S Khuvis, E Yeagle, M Cerf, and AD Mehta. Breathing above the brain stem: volitional control and attentional modulation in humans. J Neurophysiol, 119:145–159, 2018.

37. P Sörös, Y Inamoto, and RE Martin. Functional brain imaging of swallowing: an activation likelihood estimation meta-analysis. Hum Brain Mapp, 30:2426–2439, 2009.

38. H Ackermann and A Riecker. The contribution(s) of the insula to speech production: a review of the clinical and functional imaging literature. Brain Struct Funct, 214:265–272, 2010.

39. A Oh, EG Duerden, and EW Pang. The role of the insula in speech and language processing. Brain Lang, 135:96–103, 2014.

40. O Woolnough, KJ Forseth, PS Rollo, and N Tandon. Uncovering the functional anatomy of the human insula during speech. Elife, 8:e53086, 2019.

41. CJ Price. A review and synthesis of the first 20 years of PET and fMRI studies of heard speech, spoken language and reading. Neuroimage, 62:816–847, 2012.

42. JA Berken, VL Gracco, JK Chen, KE Watkins, S Baum, M Callahan, and D Klein. Neural activation in speech production and reading aloud in native and non-native languages. Neuroimage, 112:208–217, 2015.

43. JW Bohland and FH Guenther. An fMRI investigation of syllable sequence production. Neuroimage, 32:821–841, 2006.

44. A Riecker, B Brendel, W Ziegler, M Erb, and H Ackermann. The influence of syllable onset complexity and syllable frequency on speech motor control. Brain Lang, 107:102–113, 2008.

45. B Brendel, M Erb, A Riecker, W Grodd, H Ackermann, and W Ziegler. Do we have a “mental syllabary” in the brain? An fMRI study. Motor Control, 15:34–51, 2011.

46. L Ghazi-Saidi, T Dash, and AI Ansaldo. How native-like can you possibly get: fMRI evidence for processing accent. Front Hum Neurosci, 9:587, 2015.

47. A Moyer. Foreign Accent. The Phenomenon of Non-native Speech. Cambridge University Press, Cambridge, UK, 2013.

48. U Gut. Non-Native Speech: A Corpus-Based Analysis of Phonological and Phonetic Properties of L2 English and German. Peter Lang, Frankfurt, 2009.

49. M Yavas. Applied English Phonology. Blackwell, Malden, MA, USA, 2009.

